# On the design of CRISPR-based single cell molecular screens

**DOI:** 10.1101/254334

**Authors:** Andrew J. Hill, José L. McFaline-Figueroa, Lea M. Starita, Molly J. Gasperini, Kenneth A. Matreyek, Jonathan Packer, Dana Jackson, Jay Shendure, Cole Trapnell

## Abstract

Several groups recently reported coupling CRISPR/Cas9 perturbations and single cell RNA-seq as a potentially powerful approach for forward genetics. Here we demonstrate that vector designs for such screens that rely on *cis* linkage of guides and distally located barcodes suffer from swapping of intended guide-barcode associations at rates approaching 50% due to template switching during lentivirus production, greatly reducing sensitivity. We optimize a published strategy, CROP-seq, that instead uses a Pol II transcribed copy of the sgRNA sequence itself, doubling the rate at which guides are assigned to cells to 94%. We confirm this strategy performs robustly and further explore experimental best practices for CRISPR/Cas9-based single cell molecular screens.

## Introduction

Forward genetic screens in cell culture, based on RNA interference or CRISPR/Cas9, enable the functional characterization of thousands of programmed perturbations in a single experiment^1,2^. However, the options for phenotypic assays that are compatible with such screens are often limited to coarse phenotypes such as relative cell growth or survival, and moreover are uninformative with respect to the mechanism by which each positively scoring perturbation mediates its effect.

To circumvent these limitations, several groups recently reported using single cell RNA-seq (scRNA-seq)^3-5^ as a readout for CRISPR-Cas9 forward genetic screens, *i.e*. to broadly capture phenotypic effects at the molecular level. A key aspect of these methods is that the guide RNA (sgRNA) present in each single cell is identified together with its transcriptome, either by means of a Pol II transcribed barcode that is linked *in cis* to the sgRNA (CRISP-seq, Perturb-Seq, Mosaic-seq^6-9^) **(Fig. 1a)**, or alternatively by capturing the sgRNA sequence itself as part of an overlapping Pol II transcript (CROP-seq^10^) **(Fig. 1b).**

**Figure 1.**
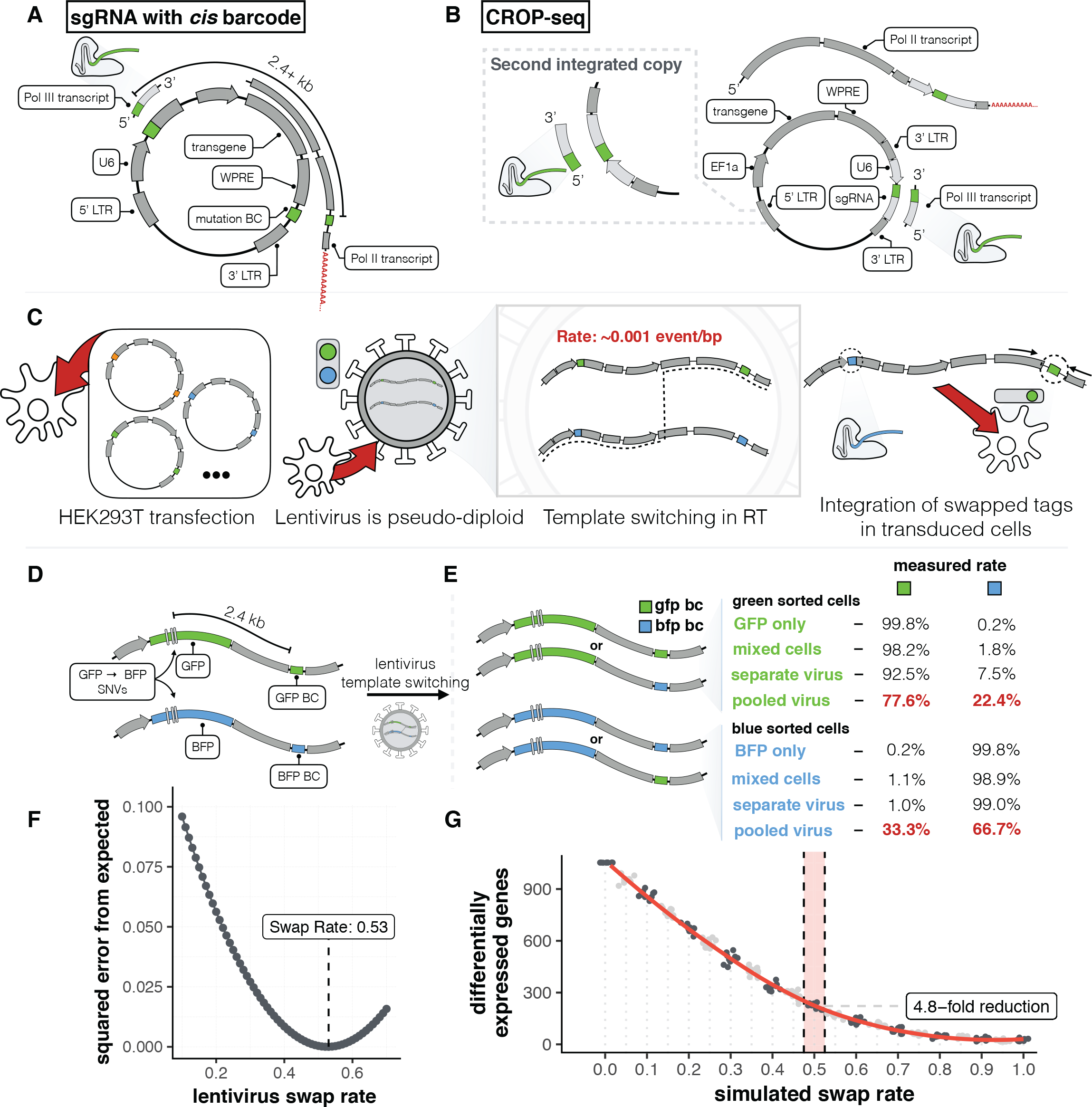
Template switching during lentiviral packaging decreases the sensitivity of designs relying on cis-pairing of sgRNAs and distal barcodes. **A)** Generalized schematic of vectors that rely on *cis* pairing of sgRNAs and barcodes. **B)** Generalized schematic of CROP-seq approach. One copy of the guide is cloned into the 3’ LTR of the vector and a second copy of the guide expression cassette is produced in the 5’ LTR during lentivirus positive strand synthesis prior to integration. **C)** Schematic of constructs developed to quantify template switching rate at 2.4 kb separation between sequences. Distinguishing bases (3 bp differences) in GFP and BFP are separated from their respective barcodes by 2.4 kb. **D-E)** Cells were transduced with GFP or BFP virus separately or a virus generated from a mix of GFP/BFP produced from individual or combined lentiviral packaging. As an additional control, cells transduced with GFP or BFP only virus were mixed prior to sorting. Cells were sorted on GFP and BFP and the percent GFP and BFP barcodes in each sample is shown as a table. Note that in a mix of two plasmids only approximately half of all chimeric products are detectable due to homozygous virions (see Methods). **F)** Plot of sum of squared errors of observed data vs. expected values at various swap rates assuming a relative proportion of 61.7% GFP+ cells as determined from FACS (see **Fig. S4** for derivation of this percentage and the supplementary methods for a detailed explanation of how the expected values are determined). **G)** Transcription factor pilot screen from Adamson *et al*., used here as a gold standard performed with arrayed lentivirus production, was subjected to simulation of progressively higher fractions of target assignment swapping to mimic the impact of template switching. Number of differentially expressed genes across the target label at FDR of 5% is plotted at each swap rate. 0.5 corresponds to the 50% swap rate determined via FACS.

In our own efforts to develop similar methods, we have encountered several important technical challenges not yet fully delineated in the literature. First, we have quantified the impact of template switching during lentiviral packaging on designs that rely on *cis* pairing of each sgRNA with a distally located barcode, and observed swapping of guide-barcode associations at a rate approaching 50%. Second, we have coupled targeted sgRNA amplification^7,8^ and the published vector of CROP-seq^10^, and shown that our improved protocol is a robust design for scRNA-seq readout of CRISPR-based forward genetic screens. Finally, we have tested an attractive alternative design to CROP-seq, but find it does not result in robust inhibition or editing.

## Results

We initially pursued a design similar to recently published methods^6-9^ in which each sgRNA was linked to a Pol II transcribed barcode located several kilobases away on the lentiviral construct **(Fig. 1a).** In our vector (pLGB-scKO), the barcode was positioned in the 3’ UTR of a blasticidin resistance transgene, such that it could be recovered by scRNA-seq methods that prime off of poly(A) tails **(Fig. S1a-b).** Guides and barcodes were paired during DNA synthesis, which facilitated pooled cloning and lentiviral delivery **(Fig. S1c).**

With this design, we sought to ask how loss-of-function (LoF) of various tumor suppressors altered the transcriptional landscape of immortalized, non-transformed breast epithelial cells^11^. Specifically, we targeted *TP53* and other tumor suppressors in the MCF10A cell line, with or without exposure to doxorubicin, which induces double-strand breaks (DSBs) and a transcriptional response to DNA damage. Cloning and lentiviral packaging was either performed individually for each targeted gene, or in a pooled fashion. In addition to scRNA-seq, we performed targeted amplification to more efficiently recover the sgRNA-linked barcodes present in each cell **(Fig. S1b; Fig. S2).**

In a first experiment in which a small number of lentiviral constructs were cloned and packaged separately for each gene (‘arrayed lentiviral production’), a substantial proportion of cells in which *TP53* was targeted had a gene expression signature consistent with a failure to activate a cell cycle checkpoint response after DNA damage, in line with *TP53*’s pivotal role in this pathway (*e.g*. lower expression of *CDKN1A* and *TP53I3;* **Fig. S3a**). However, these effects were greatly reduced when we performed a similar experiment with pooled cloning and lentiviral packaging **(Fig. S3b).** Furthermore, markedly fewer genes were differentially expressed across the targets in the pooled experiment than in the arrayed experiment **(Fig. S3c).**

Based on what is known about HIV, we reasoned that template switching during pooled packaging of lentivirus could result in the integration of constructs where the designed sgRNA-barcode pairings are partially randomized. During viral production, lentiviral plasmids are transfected into HEK293T cells at high copy number and transcribed^12^. Lentiviral virions are pseudodiploid, meaning that two viral transcripts are co-packaged within each virion^13^. The reverse transcriptase that performs negative strand synthesis has a remarkably high rate of template switching^14^, estimated as roughly 1 event per kilobase (kb)^15^. Template switching would be expected to result in the integration of chimeric products at a rate proportional to the distance between paired sequences, effectively swapping intended sgRNA-barcode associations **(Fig. 1c).** This risk was noted by Adamson et al.^7^ and Dixit et al.^8^. It was also altogether avoided by Adamson et al.^7^ through arrayed lentiviral production, but pooled lentiviral production was performed in some or all experiments of the other reports^6,8,9^. Although Sack et al.^16^ recently quantified this phenomenon at distances up to 720 bp in vectors designed for bulk selection screens, the implications of template switching at longer distances (e.g., the 2.5 kb+ separation between sgRNAs and barcodes in the pLGB-scKO, CRISP-seq, Perturb-seq, and Mosaic-seq vectors), as well as for scRNA-seq study designs specifically, remain unexplored. Given a large enough distance between the barcode and the sgRNA, *cis* linkage between the two sequences would be lost in ∼50% of integration events (an odd number of template switching events) and maintained in ∼50% (an even number of template switching events).

To quantify the extent of swapping between two distally located sequences during lentiviral packaging, we cloned BFP and GFP transgenes, which differ by three base pairs, into separate lentiviral vectors (pHAGE-GFP and pHAGE-BFP). We paired each transgene with a unique barcode, separated from the nearest unique bases in BFP/GFP by 2.4 kb **(Fig. 1d)** to approximate the 2.5 kb or greater separation between sgRNAs and barcodes in the pLGB-scKO, CRISP-seq, Perturb-Seq, and Mosaic-seq vectors^6-9^. We then transduced MCF10A cells with lentivirus generated either individually or as a pool of the two plasmids. Finally, we sorted GFP+ or BFP+ fractions of the cells with FACS, and quantified the rate of barcode swapping **(Fig. 1e; Fig. S4).** At this distance, *cis* linkage is lost at the theoretical maximum rate of 50% **(Fig. 1f; Fig. S5).** Our observations are consistent with previous estimates of template switching in HIV^15^ and recent studies in lentivirus at distances below 1 kb^16^.

In order to simulate the impact of template switching, we obtained data from a pilot experiment of Adamson *et al.^7^* generated using the Peturb-seq vector with arrayed lentiviral production, targeting several transcription factors with CRISPRi. We swapped target labels *in silico* in these data at varying rates, and evaluated the impact on power to detect differential expression. At a 50% swap rate, we observe a 4.8-fold decrease in the number of differentially expressed genes **(Fig. 1g).** This loss in power results from an effective reduction of the useful sample size for each target by twofold and contamination of each target with noise from swapped associations, thus shifting all targets towards the population average of the library.

One of the published strategies for CRISPR-based single cell molecular screens, CROP-seq^10^, does not rely on pairing of sgRNAs and barcodes. Instead, the sequence of the integrated sgRNA itself acts as a barcode, as part of an overlapping Pol II transcript. In addition, the sgRNA cassette is copied from the 3’ to 5’ LTR during positive strand synthesis **(Fig. 1b).** This copy is generated during an intramolecular priming step that does not result in intermolecular swapping at an appreciable rate^17^. A limitation of the CROP-seq method as described is that the sgRNA expressed in each cell is recovered as part of its transcriptome with limited sensitivity (∼40-60%)^10^. The roughly half of single cell transcriptomes for which the sgRNA is not identified are discarded. To improve upon this, we modified the CROP-seq protocol to include targeted amplification of the sgRNA region from mRNA libraries that have already been tagged with cellular barcodes, as in our initial pLGB-scKO design **(Fig. 2a; Fig. S6).**

**Figure 2.**
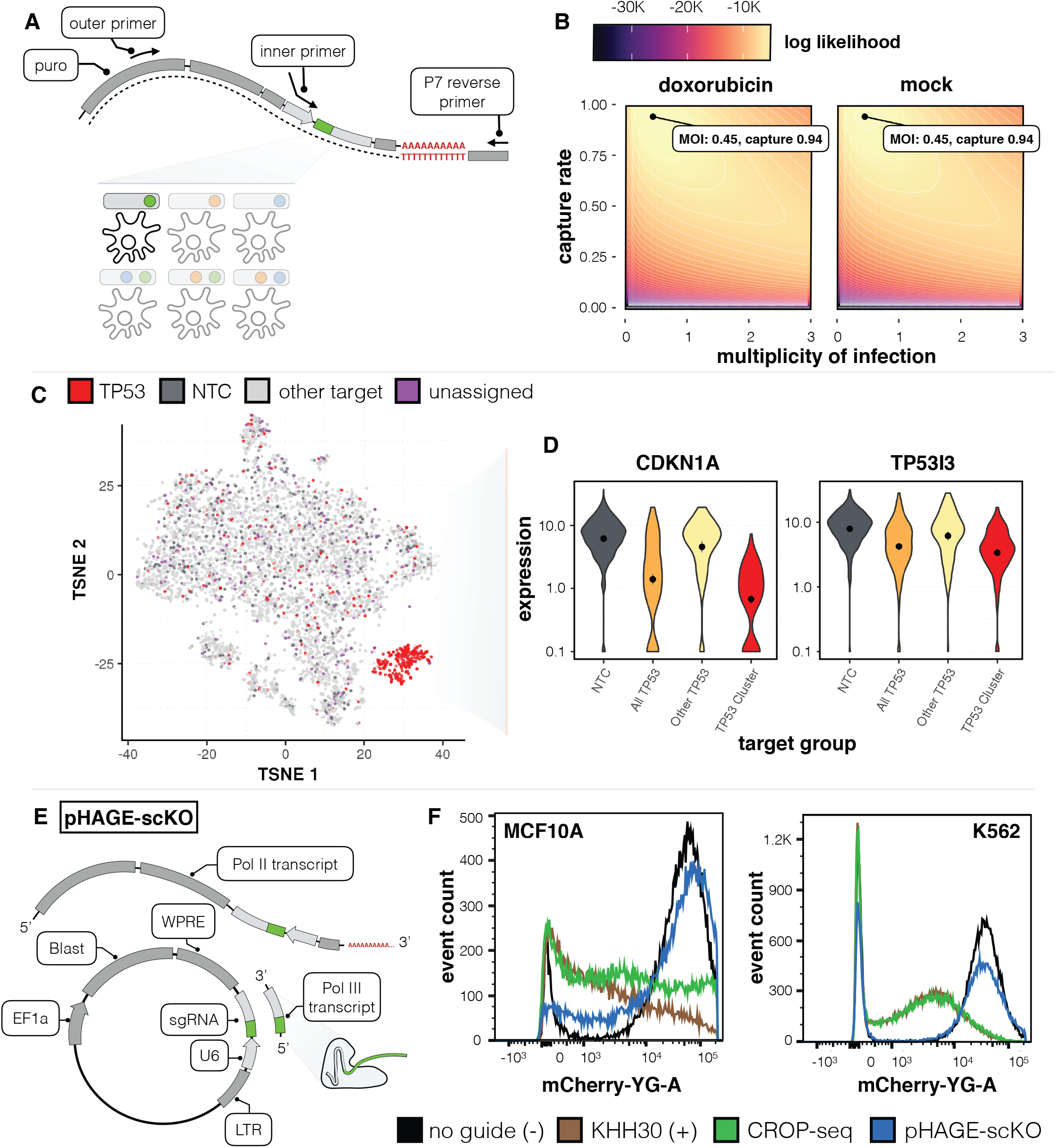
CROP-Seq screen of tumor suppressors with high capture rate by PCR enrichment, and assessment of alternate sgRNA placement within a pol II 3’UTR. **A)** Schematic of PCR enrichment of barcoded transcripts from CROP-seq samples. **B)** Determination of the most likely multiplicity of infection and capture rate of barcoded transcripts based on a generative model. **C)** tSNE embedding of a doxorubicin treated sample with colors corresponding to cells with guides to *TP53*, cells that contain non-targeting controls (NTC), cells containing guides to non-*TP53* targets, and cells that are unassigned. **D)** *CDKN1A* and *TP53I3* expression in cells expressing either non-targeting controls or guides to *TP53*. Cells with *TP53* guides are further stratified into cells inside and outside of the *TP53* enriched cluster from panel 2C. **E)** Schematic of pHAGE design with sgRNA placed upstream of the LTR. **F)** CRISPRi knock-down of mCherry in MCF10A and K562 cells not expressing a guide (- control), KHH30 (+ control), CROP-seq, and pHAGE-scKO design. All vectors have been modified to contain a CRISPRi optimized backbone.

To evaluate the effectiveness of this approach, we performed a CRISPR-mediated LoF screen of 32 tumor suppressors (6 guides per target) and 6 non-targeting control (NTC) guides in MCF10A cells with or without exposure to doxorubicin. Whereas the sgRNA was identified in the shotgun transcriptome of only 42-47% of cells, it was identified in 94% of cells with targeted amplification **(Fig. 2b).** In sharp contrast with our original pooled experiment that also targeted TP53, a tSNE embedding of doxorubicin-exposed cells from this experiment yielded a cluster that is almost entirely composed of cells containing TP53-targeting sgRNAs, highlighting the unique molecular phenotype imparted by TP53 loss when responding to DSBs **(Fig. 2c).** Specifically, the 262 cells in this cluster include 90.5% with TP53-targeting guides, 7.6% with guides targeting other genes, 0% with NTC guides, and 1.9% that were unassigned. In contrast, the remaining 5,617 cells included 3.2% with TP53-targeting guides (presumably cells in which editing failed to occur, or in which editing maintained a functional protein), 84.2% with guides targeting other genes, 7.5% with NTC guides, and 5.2% that were unassigned. Expression levels of the p53 targets *CDKN1A* and *TP53I3^18,19^* were markedly lower in the TP53-targeted cluster **(Fig. 2d)**, and 4,277 and 2,186 differentially expressed genes (FDR 5%) were identified relative to cells with NTC guides in the doxorubicin-treated and mock condition, respectively. The clean separation between the TP53-targeted cluster and other cells is presumably a consequence of: (a) the large effect of knocking out TP53; (b) our ability to recover sgRNA labels in a nearly all cells via targeted amplification; (c) the lack of sgRNA/barcode swapping, because of CROP-seq; (d) the high efficiency of CROP-seq expressed sgRNA in mediating nonhomologous end-joining.

Upon applying dimensionality reduction and clustering to cells in both the mock and doxorubicin treated conditions **(Fig. S7a-b)**, we find several other tumor suppressors whose distribution across tSNE clusters is significantly different compared to NTCs (FDR 5%), with more changes observed after doxorubicin exposure **(Fig. S7c-f).** To characterize clusters in which we observe enrichment of particular targets, we tested for target enrichment within clusters and generated average expression profiles for each enriched target-cluster pair. Gene set enrichment analysis of the most highly loaded genes in the principal components of these average expression profiles show many targets to be associated with increased proliferation and a decreased TP53/DNA damage response, with the extent of this effect being largest for TP53 **(Fig. S8).**

To further assess the impact of lentivirus template switching would have on sensitivity, we permuted target labels at varying rates within our own CROP-seq tumor suppressor screen and find a 2.9-fold reduction in the number of DEGs observed across the targets at a swap rate of 50%. Enrichment tests on our tumor suppressor screen with a 50% simulated swap rate substantially decreased the number of knockouts that display a significant change in phenotype in both mock and doxorubicin treated conditions, recovering just 4/13 *(TP53, STK11, CHEK1* and *NCOR1)* and 3/14 (TP53, *RB1*, and *ARID1B)* targets as enriched in the mock and doxorubicin conditions, respectively. Additionally, swapping simulations on the much larger (50,000 cells) unfolded protein response screen from Adamson *et al*. with arrayed lentiviral production show a 1.9- and 2.8-fold reduction at a simulated swap rate of 50% when using 25,000 and 6,000 cells, respectively **(Fig. S9).** These simulations demonstrate that the effect of barcode-sgRNA pair swapping is dependent on the number of cells captured and also on sequencing depth, number of targets examined and the effect size for those targets.

Although the CROP-seq design is not subject to sgRNA-barcode swapping, it is potentially limited by its placement of the sgRNA cassette in the LTR of the lentiviral vector, as larger intervening sequences may render the LTR non-functional^10^. To enable incorporation of longer cassettes, such as dual sgRNA designs^20-22^, we sought to place the sgRNA cassette between the WPRE and LTR. In this design (pHAGE-scKO), a second copy of this cassette would not be generated. However, the guide sequence would still contribute to overlapping Pol II and Pol III transcripts; for the former, it is positioned to ensure its incorporation into the 3’ end of the blasticidin resistance gene **(Fig. 2e).**

To evaluate this design, we compared the ability of pHAGE-scKO, CROP-seq, and a standard lentiviral sgRNA expression vector, pKHH030^23^, all containing a CRISPRi optimized backbone, to inhibit transcription via CRISPRi. We targeted the promoter of lentivirally-integrated mCherry in both MCF10A and K562 cells, which were then assayed for fluorescence via FACS. Whereas pKHH030 and CROP-seq exhibited efficient inhibition of mCherry, pHAGE-scKO had poor efficacy in both cell lines **(Fig. 2f).** Consistent with this, we observed low editing rates with our new design in MCF10A cells (88% edited with pLGB control vs. 29% edited with pHAGE-scKO). Recent studies have hinted at interference when Pol II and Pol III transcripts overlap^24,25^. We hypothesize that the observed decrease in editing and inhibition efficiency of the pHAGE-scKO design is due to the blasticidin resistance gene (Pol II promoter) inhibiting expression of the Pol III sgRNA. In contrast, CROP-seq likely maintains efficacy because the second integrated copy of the guide expression cassette (copied to the 5’ LTR during positive strand synthesis) does not overlap a Pol II transcript.

## Discussion

CRISPR-based forward genetic screens that rely on scRNA-seq to phenotype each perturbation have the potential to be extremely powerful. However, as we demonstrate here, there are important technical considerations that must be taken into account. Several published designs, as well as our own initial design (pLGB-scKO), suffer from high rates (50%) of swapped sgRNA-barcode associations, consequent to template switching across the several kilobases between the sgRNA and barcode during lentiviral production. Importantly, we do not expect that positive conclusions drawn by published studies utilizing such designs in conjunction with pooled lentivirus production^6,8,9^ are incorrect. Each of these studies examined a small number of targets and collected large datasets ranging from 25,000 to 100,000 cells per screen with ample sequencing depth, raising their baseline sensitivity. However, given the high cost of single-cell capture and sequencing and the need to expand the number of targets in such screens, our observation of ∼50% swapping of sgRNA-barcode associations with pooled lentiviral production using vectors in which the sgRNA and barcode are separated by several Kb, are highly relevant for ongoing and future studies. This loss of power may be overcome in part by filtering cells that appear inconsistent with their assigned knock-out^8^, or overcome altogether by performing cloning and lentiviral packaging separately for guides targeting each gene^7^. However, computational filtering of cells has the potential to introduce biases, and itself reduces power by discarding collected data, while performing cloning and lentiviral packaging separately for each sgRNA dramatically limits scalability.

We also explored an alternative design (pHAGE-scKO), that like CROP-seq allows sequencing of the sgRNA sequence directly, in hopes that it would facilitate the use of dual guide RNAs or other designs that require longer cassettes. However, we observe markedly reduced editing/inhibition with this design. It is plausible that methods such as programmed multiplexed guide expression cassettes^20-22,26^ could be used in conjunction with CROP-seq due to their reduced length, but it will be important to carefully validate any such designs.

As the community increasingly adopts scRNA-seq as a readout for forward genetic screens, we believe that each of these technical hurdles merit careful consideration. By coupling targeted sgRNA amplification and the published CROP-seq method^10^, we doubled the proportion of cells in which guides are assigned to cells, to 94%. The attractive features of this approach include the simplicity of the cloning protocol, its compatibility with lentiviral delivery, the high rate of recovery of sgRNA-cell associations, and no risk of template switching.

## Acknowledgements

We thank all members of the Shendure and Trapnell labs for feedback on our manuscript and helpful discussions, particularly Sanjay Srivatsan, Greg Findlay, Aaron McKenna, Riza Daza, Beth Martin, Martin Kircher, Darren Cusanovich, Xiaojie Qiu, and Vijay Ramani. We thank Professors Jesse Bloom and Douglas Fowler for discussions about lentivirus; Dr. Kyuho Han, James Ousey, and Professor Mike Bassik for experimental advice and reagents for CRISPRi experiments. AH thanks Stella the cat for support. This work was supported by the following funding: NIH DP1HG007811 and UM1HG009408 (to JS), DP2HD088158 (to CT), and the W. M. Keck Foundation (to CT and JS). AH and MG are funded by the National Science Foundation Graduate Research Fellowship. JLM is supported by the NIH Genome Training Grant (5T32HG000035) and the Cardiovascular Research Training Grant (4T32HL007828). CT is partly supported by an Alfred P. Sloan Foundation Research Fellowship. JS is an Investigator of the Howard Hughes Medical Institute.

## Data availability

Data is available on GEO via accession GSE108699 and code along with additional data are available on Github as described below. pHAGE-GFP, pHAGE-BFP, and the CROP-seq vector with the CRISPRi-optimized backbone sequence described in methods are available on Addgene as 106281, 106282, and 106280, respectively (currently pending). The CROP-seq vector with the standard backbone was used as described in Datlinger *et. al* (see methods).

## Code Availability

Code for useful scripts and analysis from the paper, more details on data access (as mentioned above), and 10X V2 enrichment PCR details are available at: https://github.com/shendurelab/singlecellkoscreens.

## Methods

### Cell Lines and Culture

MCF10A immortalized breast epithelial cells were purchased from ATCC and cultured in DMEM/F12 (Invitrogen) supplemented with 10% FBS, 1% penn-strep, 10 ng/mL EGF, 1 μg/mL hydrocortisone, 5 μg/mL insulin and 100 ng/mL cholera toxin.

### Generating an Inducible Cas9 Expressing MCF10A Cell Line

Lentivirus containing a doxycycline inducible and constitutively expressed Cas9 construct was produced by transfecting 293T cells with either pCW-Cas9 (Addgene #50661) or lentiCas9-Blast (Addgene #52962) and plasmids from the ViraPower Lentiviral Expression System (Thermo) according to manufacturer's instructions. 48 hours post transfection, virus containing supernatant was collected and cell debris removed by filtering through a 40 μm syringe filter. MCF10A cells were transduced with filtered supernatant for 48 hours and selected with 1 μg/mL puromycin (pCW-Cas9) or 10 μg/mL blasticidin (lentiCas9-Blast) for 96 hours. For cells expressing a doxycycline inducible Cas9 single cell clones of MCF10A-Cas9 cells were generated by high rate of dilution, individual clones expanded and Cas9 expression of individual clones was confirmed by immunoblotting of cells 96 hours following addition of doxycycline at 1 microgram/mL. lentiCas9-Blast cells were maintained as a polyclonal line.

pCW-Cas9 cells were used for initial arrayed and pooled screens, as well as quantification of editing rates in pHAGE-scKO vector. lentiCas9-Blast cells were used for all CROP-seq experiments.

### Initial Tagged Transcript Cloning Method

Due to our results demonstrating high rates of barcode/sgRNA swapping when using this design, we do not recommend use of this protocol.

Starting with the standard lentiGuide-puro plasmid (Addgene #52963), this vector was modified to confer blasticidin resistance, a mechanism of selection independent from the pCW-Cas9 (puro resistance) plasmid used to generate MCF10A-Cas9 cells. Puro and its EF-1A promoter were removed via a double digest with NEB SmaI (8 hours at 25 degrees C) and MLU1-HF (8 hours 25 degrees C). This product was gel purified using QiaQuick Gel Extraction kit (Qiagen). EF-1A promoter and Blasticidin, each with 20 bp homology on both ends were prepared via PCR from lentiCas9-Blast and gel purified. Fragments were assembled into digested lentiGuide-puro vector using the NEBuilder HiFi DNA Assembly kit with inserts in 2-fold molar excess and transformed into NEB C3040H E. Coli and allowed to incubate overnight at 30 degrees C. Clones were picked from plate, allowed to grow in LB+amp overnight at 30 degrees, and were purified using Qiagen Miniprep kit. Individual clones were validated via Sanger sequencing.

Lentiguide-blast was linearized using a digest with BsmB1 (Thermo) at 37 degrees for five hours followed by digestion with SalI HF (NEB) overnight and gel purification. Oligos containing both guide sequences and their corresponding barcodes were designed according to the following: tGTGGAAAGGACGAAACACC[G][guide]gttttagagctaGAAAtagcagagacgCGTCTCAgatctccctttgggccgcctccccgcg[barcode]tcgactttaagaccaatgacttaca

Where [guide] is a 20 bp guide sequence and [barcode] is an 8 bp barcode sequence uniquely paired to a particular sgRNA. Note that the [G] included prior to the guide is required for expression from a Pol III promoter. Guides that generate an extra BsmB1 restriction site when used in this design were excluded due to incompatibility with our downstream cloning strategy and only barcode sequences that did not generate additional BsmB1 restriction sites were used. RUNX1 was the only target impacted by this filter (4 guides were used instead of 6).

A library of these oligos was ordered in 96 well format as Ultramers from Integrated DNA Technologies and resuspended at equal molarity. All oligos were resuspended in water, pooled at equimolar concentrations, and amplified using a 50 microliter PCR KAPA HiFi HotStart Ready Mix PCR reaction with 1 ng of input DNA, an annealing temperature of 62 degrees, an extension time of 20 seconds, and all other parameters according to the manufacturer's recommendations. The resulting product was cleaned with a Zymo DNA Clean and Concentrator kit. The purified inserts were assembled into linearized lentiGuide-blast using the NEBuilder HiFi DNA Assembly kit and a molar excess of 1:5 vector to insert ratio. Assembled products were transformed into NEB C3040H E. Coli and grown overnight at 30 degrees in LB+amp. Product was prepared using a plasmid Miniprep kit (Qiagen).

To prepare the insert for the final reaction, a region spanning from the backbone sequence for the CRISPR sgRNA to a region towards the end of the WPRE element was amplified using the KAPA HiFi Hotstart Master Mix and purified using the Zymo Clean and Concentrator kit. The primers used in this reaction add BsmB1 cut sites that generate complementary ends in the final cloning step following digestion. This amplified fragment was ligated into PGEM-T following the kit protocol and a clone was selected via blue-white screening and validation of individual clones by Sanger sequencing. The validated construct was digested with BsmB1 (Thermo) and gel purified.

The fragment isolated from PGEM-T was then ligated into this linearized vector using a 3:1 molar excess of insert to vector using T4 DNA Ligase (New England Biolabs) and a an overnight incubation at 16 degrees C. Ligation products were transformed into NEB C3040H (stable) competent cells and grown overnight at 30 degrees in LB+amp. Plasmids were recovered using a Plasmid Miniprep kit (Qiagen).

### pHAGE Vector Cloning

The pHAGE_dsRed_IRES_zsGreen vector was modified to contain a multiple cloning site as described in *Quantification of Template Switching in Lentivirus Packaging Using FACS*. The U6-sgRNA cassette containing a 500bp filler removable by Bsmb1 digest was ordered as a gblock (Integrated DNA Technologies). Using the multiple cloning site, the U6-sgRNA cassette was added in the three-prime UTR of the zsGreen/dsRed transgene via Gibson assembly. This vector was further modified to remove the zsGreen/IRES/dsRed cassette and replace the CMV promoter with an EF1a promoter.

The vector was digested following the protocol outlined in Sanjana and Shalem *et al*. ^27^. Oligos corresponding to individual guides with homology for gibson assembly were ordered as standard DNA oligos in 96-well plate format from Integrated DNA Technologies with the following design:

[GCCTTATTTTAACTTGCTATTTCTAGCTCTAAAAC][GUIDE RC][C][GGTGTTTCGTCCTTTCCACAAGAT]

Guide RC refers to the reverse complement of the guide sequence. The entire construct may also be reverse complemented, allowing the guide sequence itself to be used rather than the reverse complement. Note that the additional C included here is required for transcription from the Pol III promoter.

All oligos were resuspended in water, pooled at equimolar concentrations, and amplified using a 50 microliter PCR KAPA HiFi HotStart Ready Mix PCR reaction with 1 ng of input DNA, an annealing temperature of 62 degrees, an extension time of 20 seconds, and all other parameters according to the manufacturer's recommendations. The following primers were used for amplification:

Forward: 5 - GCCTTATTTTAACTTGCTATTTCTAGCT - 3

Reverse: 5 - ATCTTGTGGAAAGGACGAAACA - 3

Reactions were monitored with qPCR and stopped just prior to saturation.

These reactions were cleaned with a Zymo DNA Clean and Concentrator kit and cloned into the Bsmb1 digested pHAGE vector backbone using the Clontech Infusion HD Cloning Kit. Ligations were performed using 10 fmols of vector and and a 200 fmols of double stranded oligo (1:20 molar ratio of vector to insert). Ligation products were transformed into NEB C3040H (stable) cells according to manufacturer recommendations. Transformations were diluted with 250 μL of LB and spread onto 6 LB-AMP plates and incubated at 30 degrees C for 24 hours. Colonies were then scraped into LB, a bacterial pellet was collected and plasmids recovered using a Plasmid Midiprep kit (Qiagen).

### Quantification of Template Switching in Lentivirus Packaging Using FACS

A multiple cloning site was cloned into pHAGE_dsRed_IRES_zsGreen lentiviral vector between the WPRE and 3’LTR. The multiple cloning site was assembled from annealing and extension of WPRE_MCS_insert_W and WPRE_MCS_insert_R:

WPRE_MCS_insert_W:

5- ctttgggccgcctccccgcctgggcgcgccATAACAgctagcTGATGGctcgagcc -3

WPRE_MCS_insert_R:

5- cagctgccttgtaagtcattggtcttaaaggctcgagCCATCAgctagcTGTTATgg -3

The plasmid was amplified by inverse PCR with pHAGE_WPRE_MCS_GIBS_F and R:

pHAGE_WPRE_MCS_GIBS_F

5- TGGctcgagcctttaagaccaatgacttacaaggcagctg -3

pHAGE_WPRE_MCS_GIBS_R

5- ctagcTGTTATggcgcgcccaggcggggaggcggcccaaag -3

The two fragments were cloned by Gibson Assembly. Correct clones of pHAGE_dsRed_IRES_zsGreen_WPRE_MCS were identified by Sanger sequencing and expression of the fluorescent proteins after transfection and lentiviral packaging.

To make the pHAGE EBFP or EGFP_IRES_dsRed_WPRE_MCS, pHAGE_dsRed_IRES_zsGreen_WPRE_MCS was cut with BamHI and ClaI to remove the zsGreen and IRES. The ends were blunted and re-ligated to make pHAGE_dsRed _WPRE_MCS. EGFP or EBFP (amplified with eGFP_gibsF and eGFP_IRES_GibsR) and an IRES (IRES_GibsF, IRES_GibsR) were cloned into the NotI site 5’ of the dsRed, by Gibson Assembly. EBFP was ordered as a gBlock (Integrated DNA Technologies) with 3 nucleotide changes from EGFP. Correct clones were identified by sequencing. The dsRed is not expressed in this construct.

eGFP_gibsF:

5- gccatccacgctgttttgacctccatagaagacaccggcATGGTGAGCAAGGGCGAGGAG -3

eGFP_IRES_GibsR:

5- ggatccCTACTTGTACAGCTCGTCCATGCCG -3

IRES_GibsF:

5- ATCACTCTCGGCATGGACGAGCTGTACAAGTAGggatccctcccccccccctaacgttac -3

IRES_GibsR:

5- ctccttgatgacgtcctcggaggaggccatggcggccatgtgtggccatattatcatcgtgtttttcaaagg -3

EBFP 5-

ATGGTGAGCAAGGGCGAGGAGCTGTTCACCGGGGTGGTGCCCATCCTGGTCGAGCT GGACGGCGACGTAAACGGCCACAAGTTCAGCGTGTCCGGCGAGGGCGAGGGCGATG CCACCTACGGCAAGCTGACCCTGAAGTTCATCTGCACCACCGGCAAGCTGCCCGTGC CCTGGCCCACCCTCGTGACCACCCTGACCCACGGCGTGCAGTGCTTCAGCCGCTACC CCGACCACATGAAGCAGCACGACTTCTTCAAGTCCGCCATGCCCGAAGGCTACGTCC AGGAGCGCACCATCTTCTTCAAGGACGACGGCAACTACAAGACCCGCGCCGAGGTG AAGTTCGAGGGCGACACCCTGGTGAACCGCATCGAGCTGAAGGGCATCGACTTCAA GGAGGACGGCAACATCCTGGGGCACAAGCTGGAGTACAACTTtAACAGCCACAACG TCTATATCATGGCCGACAAGCAGAAGAACGGCATCAAGGTGAACTTCAAGATCCGC CACAACATCGAGGACGGCAGCGTGCAGCTCGCCGACCACTACCAGCAGAACACCCC CATCGGCGACGGCCCCGTGCTGCTGCCCGACAACCACTACCTGAGCACCCAGTCCGC CCTGAGCAAAGACCCCAACGAGAAGCGCGATCACATGGTCCTGCTGGAGTTCGTGA CCGCCGCCGGGATCACTCTCGGCATGGACGAGCTGTACAAG -3

Fifteen nucleotide barcodes (lenti-barcode and lenti-barcode-r) were then cloned into the multiple cloning site between the WPRE and 3’LTR for both the EBFP and EGFP constructs by Gibson Assembly. Single clones were prepared and the barcode identified by Sanger sequencing.

lenti-barcode:

5- atctccctttgggccgcctccccgcctgggGGATCCAGNNNNNNNNNNNNNNNtcgagcctttaagaccaatgactt acaagg -3

lenti-barcode-r:

5- CCTTGTAAGTCATTGGTCTTAAAGGCTCGA -3

Lentivirus was packaged by transfection of barcoded EGFP or EBFP constructs either alone or in an equimolar mix along with helper plasmids (pHDM-Hgpm2, pHDM-Tatlb, pRC-CMVRev1b and pHDM-VSV-G) into HEK293T cells using Lipofectamine 2000 (Invitrogen). Viral supernatant was collected after 48 hours, spun to remove debris, snap frozen in liquid nitrogen and stored at -80°C. To titer the packaged lentiviruses, they were thawed on ice and added to MCF10A cells with media containing 8 micrograms/ml polybrene, andthe frequency of transduced cells 48 hours post-transduction was determined by flow cytometry.

To sort blue+ and green+ populations, 400,000 of MCF10A Δ*TP53* cells (Horizon Discovery) in 5 ml media plus 8 micrograms/ml polybrene were transduced at a MOI ∼0.1, with either of the EGFP or EBFP expressing viruses that had been packaged singly, a mix of the EGFP and EBFP expressing viruses that had been packaged singly or the EGFP and EBFP expressing viruses that had been packaged together. The cells were then cultured for four weeks to avoid residual plasmid contamination following transduction. An equal number of cells transduced with EGFP and EBFP virus were mixed to determine the rate of contamination resulting from FACS error. The mixed cells along with others were sorted for blue+ or green+ populations using a FACS Aria II (Becton Dickinson) that had been compensated for the overlap between the EBFP and EGFP emission spectra. The genomic DNA was harvested from each population using the Qiagen DNeasy kit. The barcodes were amplified from 2-36 ng of genomic DNA in 50 ul Robust polymerase (Kapa) reactions with primers bwds_p5_WPRE_BC_F and bwds_next_WPRE_BC_R.

bwds_next_WPRE_BC_R:

GGCTCGGAGATGTGTATAAGAGACAG

5- gaaatcatcgtcctttccttggct -3

bwds__p5_WPRE_BC_F:

5- AATGATACGGCGACCACCGAGAgcgccgatgccttgtaagtcattggtcttaaaggctc -3

Reactions were removed from the thermocycler just prior to saturation (27-30 cycles). The PCR products were purified with Ampure (Agilent) and P7 index sequences were added by an additional six cycles of PCR. PCR products were purified, quantified, pooled and single-end sequenced on an Illumina Nextseq500 with Read1 primer bwds_WPRE_bc_seqF and standard Illumina i7 primers.

bwds_WPRE_bc_seqF:

5- GCG CCG ATG CCT TGT AAG TCA TTG GTC TTA AAG GCT CGA -3

### Analysis of FACS Data from pHAGE-GFP and pHAGE-BFP Experiments

The background percentage of contaminating barcodes in the BFP/GFP sorted cells from the mixed cells control was first subtracted from the numbers obtained for the pooled virus samples. The fraction of GFP cells, as determined from FACS gating, was fixed and the expected fraction of barcode contamination in the BFP and GFP given this fixed fraction of GFP cells was simulated. Note that the expected contamination of green barcodes in the BFP sorted cells is simply the template switching rate multiplied by the fraction of green cells. The expected rate of contamination of BFP barcodes in the GFP sorted cells is simply the template switching rate multiplied by the BFP cell fraction (1 - GFP cell fraction). The sum of the squared error between the observed and expected values for these to rates of contamination was calculated for a range of different lentivirus swap rates and the minimal value was taken to be the most likely swap rate given the observed data.

Note that, unlike in a large library of plasmids, in a mix of two plasmids, only half of all chimeric products formed will be detectable as many virions will be homozygous (contain the same construct, and thus chimeric products are identical to the original). To give an analogous example, in a barnyard experiment for a single-cell assay, mouse-mouse or human-human multiplets cannot be detected and thus estimated rates of ‘doublets’ have to be adjusted accordingly. When the plasmids are equimolar and the swap rate is 50%, for example, one would expect to observe a 75% rate of the intended barcode and a 25% rate of the unintended barcode.

This ratio will change according to the molar concentration of the two plasmids. In **Fig. 1f**, we assume that the pool was composed of 61.7% GFP plasmid, corresponding to the fraction of GFP+ cells relative to the total number of GFP+ and BFP+ cells -- 4.59 / (4.59 + 2.85) or 61.7% as explained in **Fig. S4.** This analysis was also performed without fixing the fraction of GFP+ cells to the value measured by FACS to ensure results were concordant between the two methods **(Fig. S5).** The minimum sum of squared error over the grid of simulated lentivirus swap rate and fraction of GFP cells were taken to be the most likely set of parameter values.

### CRISPRi Experiment

K562 expressing dCas9-BFP-KRAB (gift of the Bassik lab, Addgene #46911) and MCF10A expressing dCas9-BFP-KRAB (made by transduction with lenti_UCOE_EF1-dCas9-BFP-KRAB, plamid available on Addgene soon; see https://weissmanlab.ucsf.edu/CRISPR/CRISPRiacelllineprimer.pdf) were transduced with lenti-mCherry under control of a CAG-promoter (pCAG_mCherry pKH143, gift of the Bassik lab, unpublished), and sorted such that the resulting population is enriched for mCherry expression.

A spacer targeting the CAG-promoter was cloned into the KHH030 (Addgene #89358), CROP-seq, and pHAGE sgRNA expression vectors. The CROP and pHAGE were modified by Q5-Site Directed Mutagenesis (New England BioLabs) to use the previously described sgRNA-(F+E)-combined optimized backbone ^28^. The CRISPRi mCherry+ K562 and MCF10A cells were transduced with the CAG-targeting sgRNA, and again assayed for mCherry.

All virus for the CRISPRi experiments were made by the Co-operative Center for Excellence in Hematology Vector Production core. All sorting was performed on a FACS Aria II (Becton Dickinson).

### Editing Rate Experiment for pHAGE-scKO

To confirm that our pHAGE-scKO vector exhibited reduced editing efficiency in addition to reduced inhibition efficiency via CRISPRi, we performed editing with a guide to TP53 from our screen (GAGCGCTGCTCAGATAGCGA) in both lentiGuide-Blast and pHAGE-scKO using our pCW-Cas9 MCF10A cells. Cells were passaged for 18 days post-induction of Cas9 expression with dox and gDNA was harvested using Qiagen DNeasy kit and amplified using primers CTAAATGGCTGTGAGAGAGCTCAGCCACACGCAAATTTCCTTCC and ACTTTATCAATCTCGCTCCAAACCCCCTGCCCTCAACAAGATGT. These were then amplified using KAPA HiFi Hotstart Ready Mix (KAPA) using the following primers to generate final indexed sequencing libraries: AATGATACGGCGACCACCGAGATCTACACacgtaggcCTAAATGGCTGTGAGAGAGCTC AG

CAAGCAGAAGACGGCATACGAGAT[INDEX]gaccgtcggcACTTTATCAATCTCGCTCCA AACC

These reads were then processed using the method described in McKenna and Findlay *et al. ^29^* Briefly, low quality bases are trimmed using Trimmomatic, reads are merged using Flash, aligned to the reference of the locus surrounding the guide site using needle, and unique genotypes are quantified. The wild-type genotype fraction was taken to be the proportion of unedited alleles. We did not use UMIs in this experiment. The lack of UMIs may overestimate the editing rate in all samples to some extent due to amplification bias.

### KO Experiments

For all screens, each plasmid library was transfected along with plasmids provided with the ViraPower Lentiviral Expression into 293T cells. At 48 and 72 hours post transfection, viral containing supernatant were collected, filtered using a 40 μm steriflip filtration system (EMD Millipore). For arrayed experiments, individual plasmids were transfected and viruses produced as described above. For pHAGE-scKO and arrayed/pooled pLGB-scKO vector experiments, virus concentrated using Peg-it virus concentration solution (SBI). Viral titer of the concentrated lentiviral library was determined by transduction of MCF10A-Cas9 cells for 48 hours at several viral dilutions, splitting cells into replica plates, and subjecting replica plate to blasticidin. Percent control growth was used to assess MOI. MCF10A-Cas9 cells with estimated MOIs of 0.3 were carried forward for further experiments.

For pHAGE-scKO and arrayed/pooled pLGB-scKO vector experiments, media was switched to 1 microgram/mL doxycycline to induce expression of Cas9 in pCW-Cas9 cells. LentiCas9-Blast cells were used for CROP-seq experiments, which do not require induction of Cas9 expression. Editing was allowed take place for 14 days for arrayed and pooled pLGB-scKO and 21 days for pHAGE-scKO and CROP-Seq experiments. Media was changed every 48 hours and cells were cultured every 96 hours. For the first half of editing, cells were cultured in the presence of 5 μg/mL blasticidin and 0.5 μg/mL puromycin to ensure high sgRNA and Cas9 expression.

### Doxorubicin Treatment

After editing, MCF10a cells were seeded in 10 cm plates plates at 1 x 10^6^ cells per well, allowed to attach overnight and media replaced with MCF10A media alone (mock) or MCF10A media containing 500 (arrayed and pooled pLGB-scKO experiments) or 100 nM (pHAGE-scKO and CROP-Seq experiments; we ultimately decided that this lower dose was more appropriate and likely to provide more robust signal) doxorubicin prepared from a 500 μM stock of doxorubicin (Sigma) in water. 24 hours after drug exposure untreated and doxorubicin treated cells were harvested by trypsinization, washed with PBS and used for downstream assays.

### Single-Cell RNA-sequencing

Cells were captured using one lane of a 10X Chromium device per sample using 10X V1 Single Cell 3’ Solution reagents (10X Genomics). Approximately 4000-7000 cells were captured per lane for each condition. Protocols were performed according to manufacturer recommendations, holding 10-30 ng of full length cDNA out of downstream shearing and library prep steps in order to provide material for barcode enrichment PCR.

Final libraries were sequenced on NextSeq 500/550. 10X V1 samples were sequenced using the following read configuration on 75 cycle High Output kits: R1: 64, R2: 5, I1: 14, I2: 8

Our initial arrayed and pooled doxorubicin treated samples using pLGB-scKO were aggregated using cellranger aggregate to normalize the average number of mapped reads per cell. This yields an average of 37,732 reads per cell, 2263 median genes per cell, and a median of 8279 UMIs per cell.

Our CROP-seq mock sample was sequenced to an average depth of 120,797 raw reads per cell in 6598 cells. A median of 4619 genes per cell were detected and a median UMI count of 22,495 per cell. Our CROP-seq doxorubicin treated sample was sequenced to an average depth of 123,445 raw reads per cell in 6283 cells. A median of 3500 genes per cell were detected and we observed a median UMI count of 15,324 per cell. At this depth the average duplication rate is approximately 78%.

### Enrichment PCR

For all experiments a hemi-nested PCR starting from 5 ng of full length cDNA was used to enrich for the barcodes that assign a target to each cell. All PCR reactions were performed with a P7 reverse primer (as introduced by the 10X Chromium V1 oligo DT RT primer). For pHAGE-scKO and pLGB-scKO, the first PCR was performed with

5- TCCTGGGATCAAAGCCATAGT -3

and for CROP-Seq with

5- TTTCCCATGATTCCTTCATATTTGC -3

as the forward primer, priming to the blasticidin transcript with no non-templated sequence for pHAGE-scKO and pLGB-scKO, and to part of the U6 promoter in CROP-seq. For pLGB-scKO the second PCR was performed with

5- TCGTCGGCAGCGTCAGATGTGTATAAGAGACAGGACGAGTCGGATCTCCCTT -3

for pHAGE-scKO with

5-

TCGTCGGCAGCGTCAGATGTGTATAAGAGACAGAACGGACTAGCCTTATTTTAACTTG -3

and for CROP-Seq with

5- TCGTCGGCAGCGTCAGATGTGTATAAGAGACAGcTTGTGGAAAGGACGAAACAC -3

as the forward primer, priming adjacent to the barcode/guide sequence in each design and adding the standard Nextera R1 primer. Samples were then indexed in a final PCR using standard Nextera P5 index primers of the form:

5- AATGATACGGCGACCACCGAGATCTACAC[10bp Index]TCGTCGGCAGCGTC -3

Each PCR was cleaned with a 1.0X Ampure XP cleanup and one microliter of a 1:5 dilution of the first PCR was carried forward and a 1:25 dilution of the second PCR was carried into the final PCR reaction. PCRs were monitored by qPCR and stopped just prior to reaching saturation to avoid overamplification. The final PCR was run on a Bioanalyzer to confirm expected product size.

### Digital Gene Expression Quantification

Sequencing data from each sample was processed using cellranger 1.3.1 to generate sparse matrices of UMI counts for each gene across all cells in the experiment.

Each lane of cells was processed independently using cellranger count, aggregating data from multiple sequencing runs. For the comparison between arrayed and pooled screens, cellranger aggregate was used to downsample data from each screen to an equal average number of mapped reads.

### Assigning Cell Genotypes

Barcode enrichment libraries were separately indexed and sequenced as spike-ins alongside the whole transcriptome scRNA-seq libraries. Final UMI and cell barcode assignments were made for each read by processing these samples with cellranger 1.3.1 as was done for the whole transcriptome libraries.

A whitelist of guide or target barcode sequences was constructed using all guides or target barcodes in the library. For each read in the position sorted BAM file output by cellranger 1.3.1, the final cellular barcode and UMI are extracted. If either of these fields is not populated, indicating low sequencing quality for the cell barcode or UMI read, the read is ignored. Using the cDNA read, we attempt to find a perfect match for the sequence immediately preceding the guide or barcode (GTGGAAAGGACGAAACACCG for CROP-seq and CGCCTCCCGCG for pLGB-scKO). If a perfect match is not found, we attempt to locate the sequence in an error-tolerant manner using a striped Smith-Watterman alignment, where alignments must span a length no more than 2bp shorter than the search sequence. If a match or alignment is found, the guide or barcode sequence is extracted. If the extracted sequence does not perfectly match a whitelist sequence, we search for a matching whitelist sequence within an edit distance of half the minimum edit distance between any pair of guides or barcodes in the library (rounded down). If no match is found, the molecule is ultimately discarded. Matches to the whitelist are tracked for each cell.

We also remove likely chimeric sequences using the approach outlined in Dixit ^30^. Briefly, within each cell, we first calculate the number of times a given UMI is observed with each observed guide assignment. We then divide these counts by the total instances of the respective UMI across all observed guide assignments within that cell. For UMI-guide assignment combinations where this fraction is less than 20%, we do not count the UMI towards the final observed guide assignment counts. While this has some impact on the raw data, we find the benefits to be modest, in contrast to results reported in Dixit *et al*. ^8^.

To make a set of final assignments, we take all whitelist sequences with over 10 reads and account for over 7.5% of the whitelist reads assigned to a given cell, where multiple sequences can be assigned to each cell. Whitelist sequences and their corresponding target genes are assigned to each cell. Finally, this set of assignments is merged with the filtered gene expression matrices output by cellranger such that only assignments to the set of high quality cells appear in the final dataset.

Note that when processing CROP-seq data without PCR enrichment, we lowered the requirement for reads supporting a given guide to 3 to account for the decreased coverage of these transcripts.

### Estimation of MOI and Capture Rate

The most likely multiplicity of infection and capture rate given the distribution of guide counts per cell were estimated using the generative model described in Dixit *et al*. ^8^. Briefly, a log likelihood is calculated using a zero-truncated poisson (represents the multiplicity of infection after selection for cells that harbor a lentiviral construct) convolved with a binomial (represents the incomplete capture of transcripts containing guide sequences from cells). This model is used to to calculate log-likelihood values for a range of MOI values and capture rate values (rate of observing a guide in a cell given that the guide is present). The maximum log-likelihood is taken to be the MOI and capture rate of the experiment.

### Removing Low Quality Cells

Despite using the filtered set of cells provided by cellranger to exclude cell barcodes with low UMI counts, we consistently observed a cluster of cells with much lower UMI counts on average than the rest of the dataset when performing dimensionality reduction. To avoid including these cells in downstream analysis, we perform a simple procedure to remove any cluster with low average UMI counts.

First, we perform PCA on the matrix of all cells and genes expressed in at least 50 cells for each condition to reduce to 12 principal components. We then reduce to a two dimensional space using tSNE. Next we perform density peak clustering in the two dimensional space using default parameters. For each cluster, we calculate the average size factor over the cluster as calculated using estimateSizeFactors in monocle2 ^31^. We observed that filtering out clusters of cells with an average size factor of -0.85 or lower readily distinguished the low quality cluster of cells. All cells contained in these clusters were removed from downstream analysis. PCA and tSNE were performed using the monocle2 function reduceDimension with default parameters and the tSNE option. Density peak clustering was performed using the monocle2 function clusterCells.

### Simulating Loss in Power from Barcode Swapping

Assignments were permuted for a fraction of cells ranging from 0 to 100% and kept fixed for the remaining fraction of cells. The monocle2 function differentialGeneTest was used to test for genes differentially expressed across the target assigned to each cell (only testing genes detectably expressed in at least 50 cells). The number of genes with a qvalue of 0.05 or lower was counted. This was performed over 10 different resamplings for each rate of swapping to obtain a distribution for each swap rate.

The average fold-reduction in DEGs resulting from a 50% swap rate was taken to be the approximate fold change in power resulting from template switching in our original design.

For the simulation performed on our own data, cells with a single target assignment from 100nm doxorubicin treated cells in our CROP-seq experiment were taken as the starting set of cells.

For the simulation on data from Adamson *et al*., processed data was obtained from GEO (GSE90546). Assignment of cells to targets were used as provided on GEO and only cells that were noted as having high quality assignment of the assigned target and noted as being a single cell were used in downstream analysis. Due to the large number of cells (50,000+) in the UPR experiment from this study and the large number of differential tests required for these simulations, the number of cells assigned to each target was downsampled by 2-fold to reduce runtime. We also performed tests on a dataset further downsampled to approximately 6,000 cells to illustrate the relationship between the initial power of a screen and the impact of simulated target swapping on sensitivity.

### tSNE Embedding Demonstrating TP53 Enriched Cluster

Scaling, PCA, and tSNE on PCA results were performed with the monocle2 function reduceDimension. 20 dimensions from PCA were carried into tSNE which was performed to two dimensions using default parameters. All cells (except those excluded in the low size factor cluster) including cells with guides to multiple targets and no assigned target were included in dimensionality reduction for this plot. Percentages of cells with guides to TP53 and ARID1B were calculated including guides that contain guides to multiple targets (all cells with TP53 guides were counted as TP53 cells and all cells without TP53 but with one or more guides to ARID1B were counted as ARID1B cells for the purposes of calculating reported percentages).

### Enrichment of Tumor Suppressors in Specific Molecular States

Only cells containing a guide to a single target were considered in all enrichment testing. A Chi squared test was used to determine whether the distribution of individual sgRNAs and targets in tSNE space was significantly different from non-targeting controls at an FDR cutoff of 5%. Targets which did not pass this test and did not have any individual sgRNA pass the test were excluded from the subsequent enrichment tests. For each sgRNA of the remaining targets, we sought to estimate the functional editing rate (probability of a cell having a true LoF given that it received that sgRNA), but such estimates would be confounded if one accounts for the possibility of edits that cause LoF for the target gene but have incomplete penetrance on the cellular phenotype. Therefore we used an expectation maximization approach to estimate the functional edit rate of each sgRNA relative to the unknown functional edit rate of the most efficient sgRNA for a given target.

The t-SNE cluster distribution of all cells in which a given sgRNA was detected was modeled as a mixture of the t-SNE cluster distribution of cells with a functional edit for the sgRNA's target gene and the t-SNE cluster distribution of non-targeting controls, where the mixing parameter is the relative functional edit rate for that sgRNA. In the expectation step, the t-SNE cluster distribution of cells with a functional edit for the target is estimated as the weighted average of the empirical t-SNE cluster distributions of each sgRNA for the target, weighted by the current estimates of the relative functional edit rate of the sgRNAs. In the maximization step, the relative functional edit rate of each sgRNA for the target is estimated as that which maximizes the likelihood of the observed t-SNE cluster distribution for cells receiving that sgRNA under the multinomial mixture model.

After estimating the relative functional edit rate for each sgRNA, a weighted contingency table was constructed where the rows are targets, the columns are t-SNE clusters, and the values are weighted cell counts, where a cell's weight is proportional to the relative functional edit rate for the sgRNA it received. Fractional values were rounded down. Fisher's exact test was applied to this weighted contingency table to test for enrichment of targets amongst t-SNE clusters. Targets were defined as enriched at an FDR of 10%. Chi square and Fisher's exact test were performed using R functions chisq.test and fisher.test, respectively.

### Principal component and gene set enrichment analysis

Pairwise differential gene expression analysis was performed between enriched target cells and non-targeting controls for cells in all significant enriched target-cluster pairs from our enrichment testing. The union of all differentially expressed genes across targets (FDR 5%) was used to perform principal component analysis. Gene set enrichment analysis was performed on genes that had the top positive and negative loadings for principal component 1 (less than -0.02 or greater than 0.02). Gene set enrichment analysis was performed using the piano R package and the hallmarks gene set from MSigDB. Gene sets were defined as enriched at an FDR cutoff of 1%. PCA was performed using the prcomp function in R, differential expression analysis was done using the monocle2 function differentialGeneTest. The hallmarks gene set collection GMT file was downloaded from the MSiGDB.

## Supplementary Figure Legends

**Figure S1.**
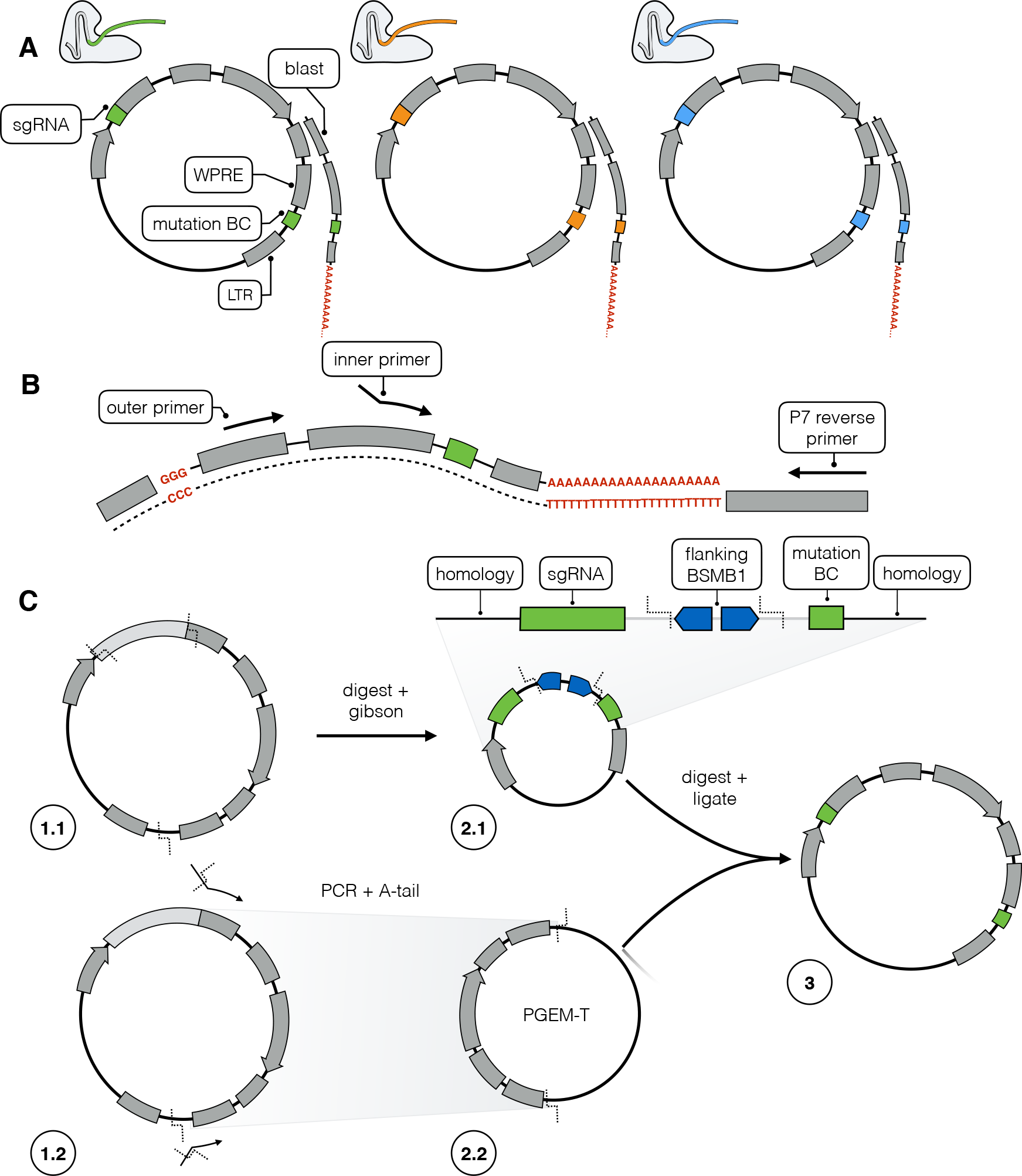
Diagram of cloning protocol and barcoded transcript enrichment strategy relying on *cis* pairing of sgRNAs and barcodes (pLGB-scKO). **A)** Schematic of our final vector relying on *cis* pairing of an sgRNA and a distal barcode. **B)** Strategy for PCR enrichment of barcoded transcripts from single-cell RNA-seq data. **C)** Pooled cloning protocol. In 1.1 we start with pLentiguideBlast and digest near the final locations of the sgRNA and paired barcode. In 2.1 an engineered library of oligos containing programmed pairs of sgRNAs and corresponding barcodes are inserted into the digested vector. In 1.2 a portion of pLentiguideBlast is amplified. In 2.2 this fragment is cloned into PGEM-T. Finally, in step 3 vectors resulting from 2.1 and 2.2 are digested with Bsmb1 and the insert from 2.2 is ligated into the backbone in 2.1 to produce the final library of sgRNAs and paired barcodes.

**Figure S2.**
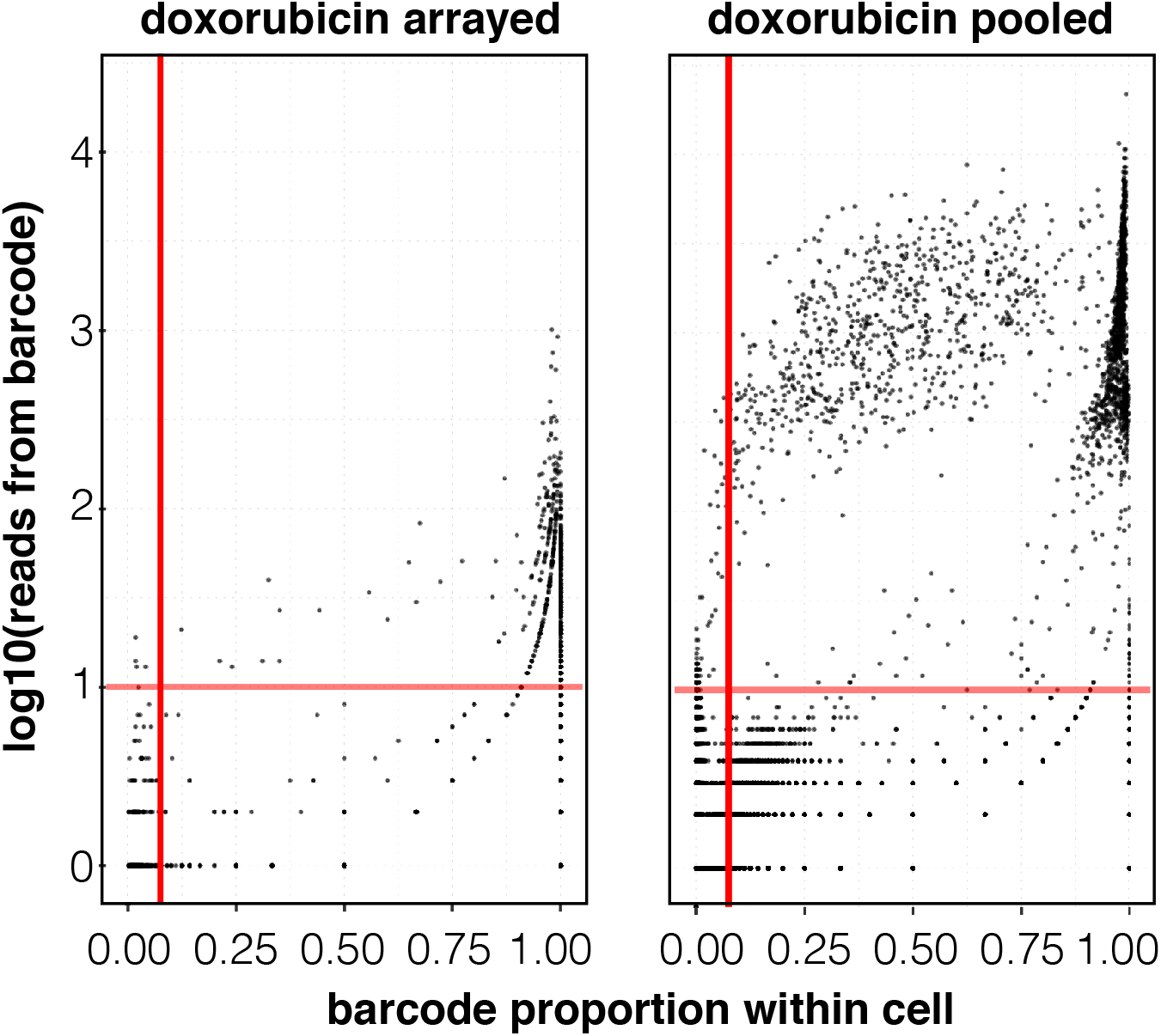
Barcoded transcript enrichment quality control for arrayed and pooled pLGB-scKO experiments. Each dot represents a barcode sequence observed in a given cell. Plot of reads for a given barcode against the proportion of all barcode reads observed in a given cell for every barcode/cell pair. Red lines indicate the lower-bounds used to distinguish noise from true barcode observations (10 reads and 0.075 proportion within cell). All barcodes observed above the red lines are assigned to their respective cells. Left, doxorubicin treated sample from arrayed experiment. Right, Doxorubicin treated sample from pooled experiment.

**Figure S3.**
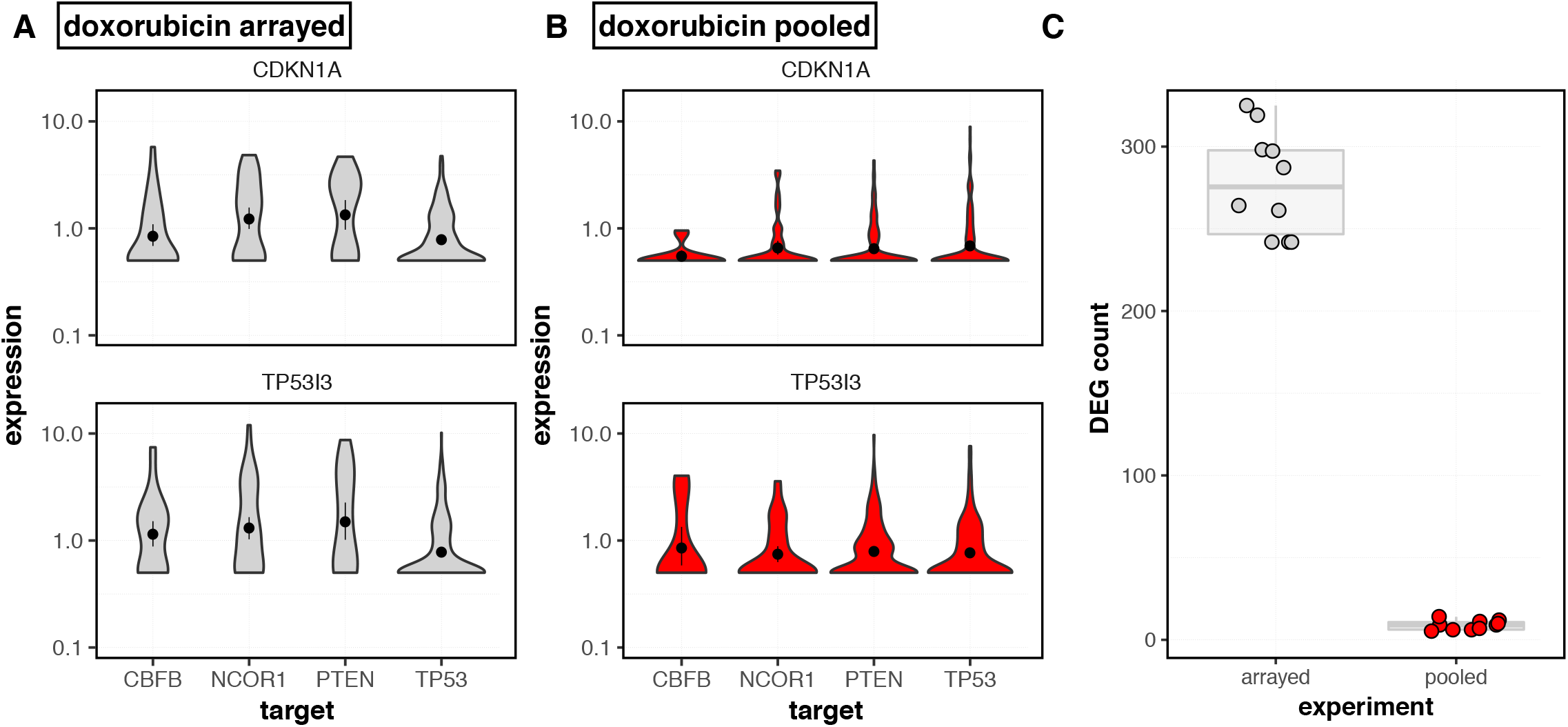
Comparison of a screen performed with arrayed and pooled lentivirus production using a vector that relies on *cis* pairing of sgRNAs and barcodes. Experiments were performed at different times but under the same conditions. The arrayed experiment was performed as a pilot experiment with 4 targets and observed an overall low rate of cells with detected barcodes. The pooled experiment was performed afterwards with 10 targets and a set of non-targeting controls and we observed a high proportion of cells with detected barcodes and good coverage of the library. To compare these experiments, only the four overlapping targets were considered and the number of cells containing an sgRNA to each target and sequencing depth were matched between samples to control for power differences. **A)** Size-factor normalized *CDKN1A* and *TP53I3* expression across *TP53* and the three other targets in arrayed screen. **B)** *CDKN1A* and *TP53I3* expression across *TP53* and three other targets in pooled screen that overlap with the arrayed screen. **C)** Comparison of the number of differentially expressed genes detected at an FDR of 5% for arrayed across the target label in the arrayed and pooled experiments.

**Figure S4.**
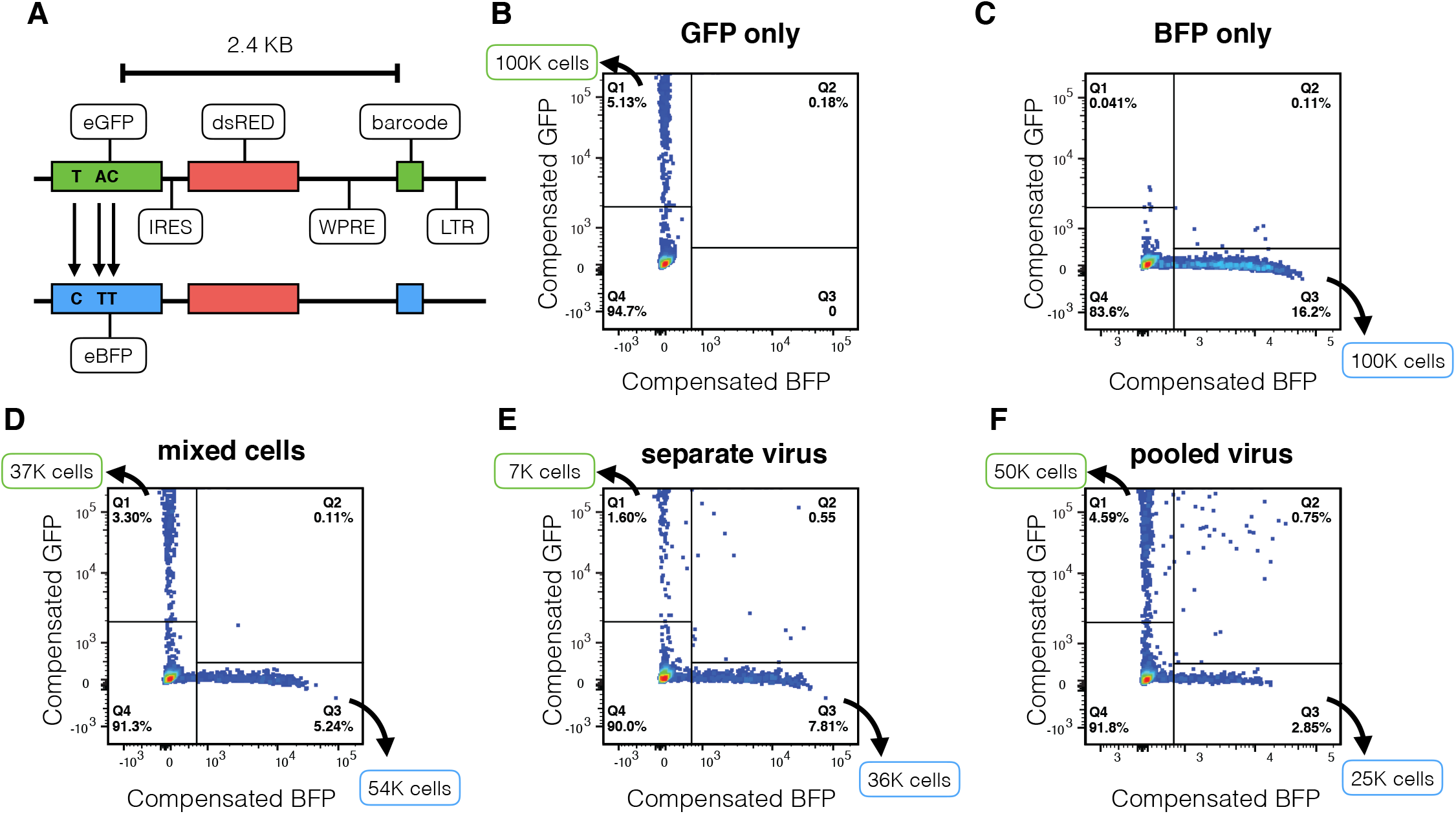
Design and sorting of GFP and BFP positive fractions in lentivirus barcode swapping experiment. **A)** Schematic of vectors (pHAGE-GFP and pHAGE-BFP) designed to quantify template switching rate at 2.4 kb using a FACS readout. FACS plots are shown for sorted cells in samples corresponding to **B)** GFP only transduced cells **C)** BFP only transduced cells **D)** GFP and BFP only transduced cells mixed just prior to FACS as a control **E)** cells transduced with BFP and GFP virus that was generated separately but pooled prior to transduction **F)** cells transduced with BFP and GFP virus that was generated from pooled plasmids. The fraction of green plasmids assumed in the determination of lentivirus swap rate from FACS experiments is taken as the fraction of GFP+ cells relative to the total GFP+ and BFP+ cells from this sort (4.59 / (4.59 + 2.85) or 61.7%). This accounts for the fact that plasmids were likely not completely equimolar. The approximate number of total cells cells sorted in each fraction is indicated along the appropriate axes on each plot.

**Figure S5.**
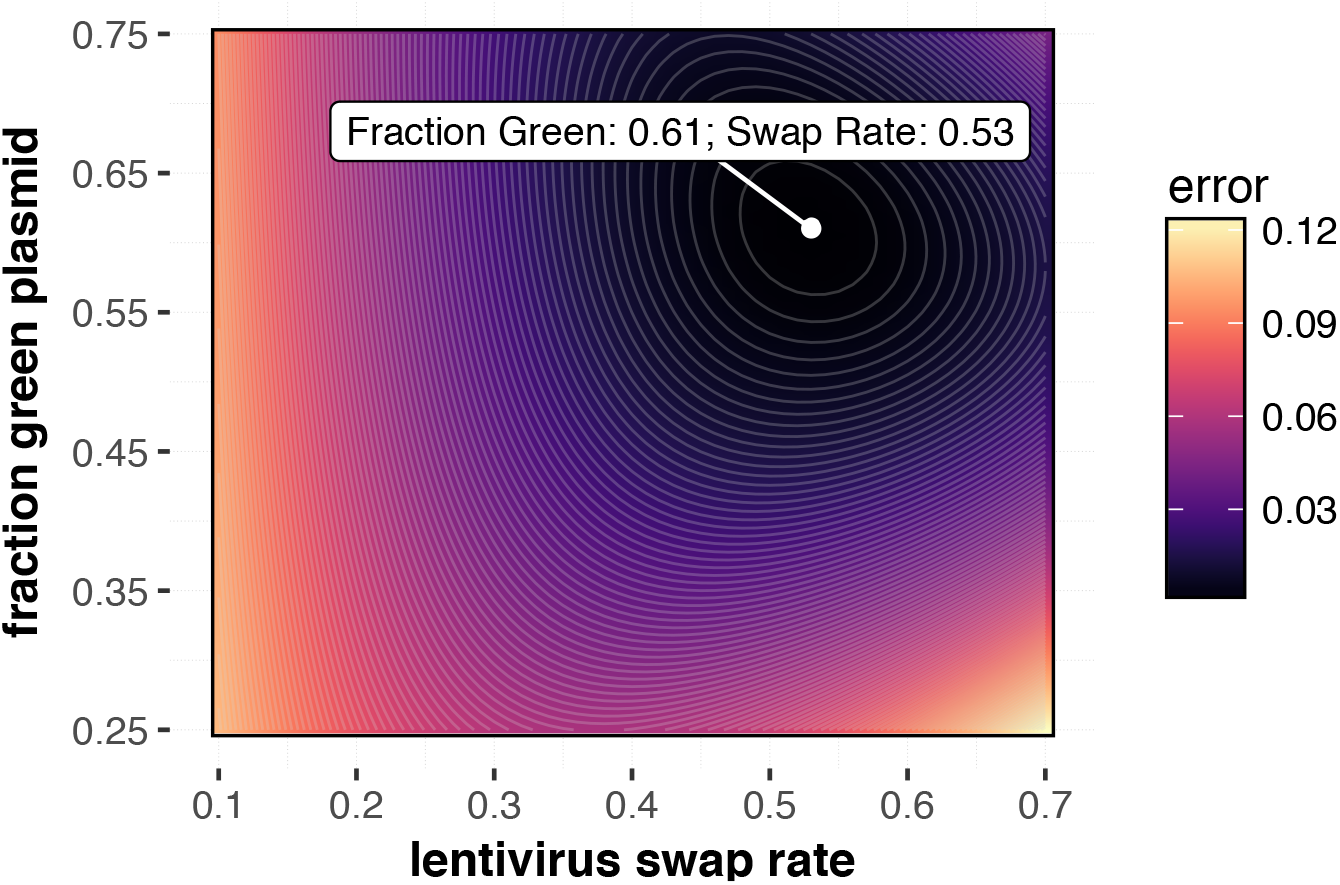
Simulation of concordance between observed and expected data obtained from FACS experiment in **Fig. S4** to quantify template switching rate at 2.4 kb separation between paired sequences. **Fig. 1F** assumed a fraction of 0.617 of GFP plasmid in the original green plasmid / blue plasmid mix as determined from FACS in **Fig. S5.** In this figure, both the fraction of GFP plasmid and lentivirus swap rate are varied to obtain the set of parameters that best fit the collected data. The sum of the squared error between expected and observed values from FACS given each combination of parameters is shown.

**Figure S6.**
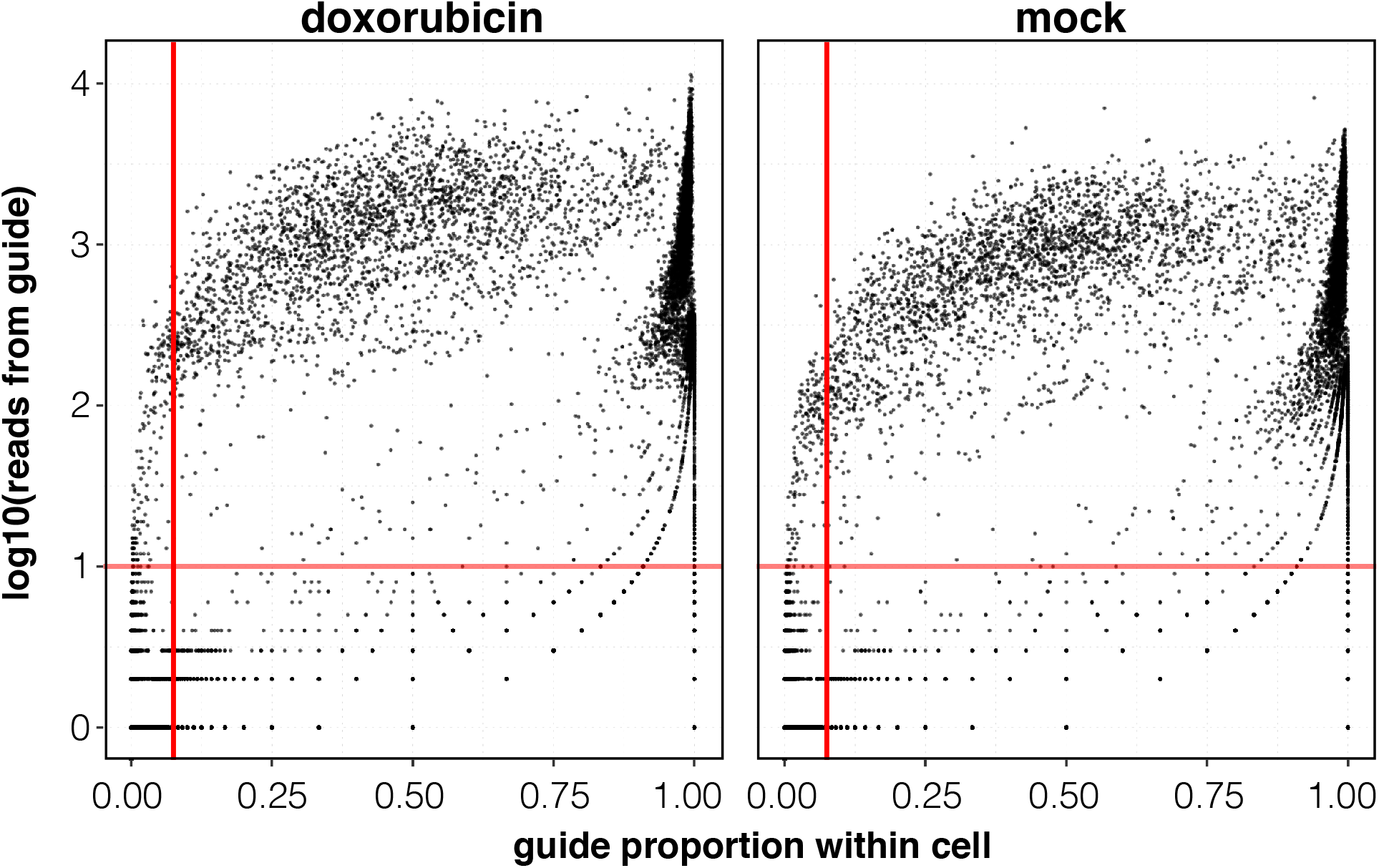
Guide transcript enrichment quality control plot for tumor suppressor knock-out screen performed with CROP-seq. Each dot represents a guide sequence observed in a given cell. Plot of reads for a given guide against the proportion of all guide reads observed in a given cell for every barcode/cell pair. Red lines indicate the lower-bounds used to distinguish noise from true guide observations (10 reads and 0.075 proportion within cell). All guide observed above the red lines are assigned to their respective cells. Left, Doxorubicin treated sample from CROP-seq experiment. Right, Mock sample from CROP-seq experiment.

**Figure S7.**
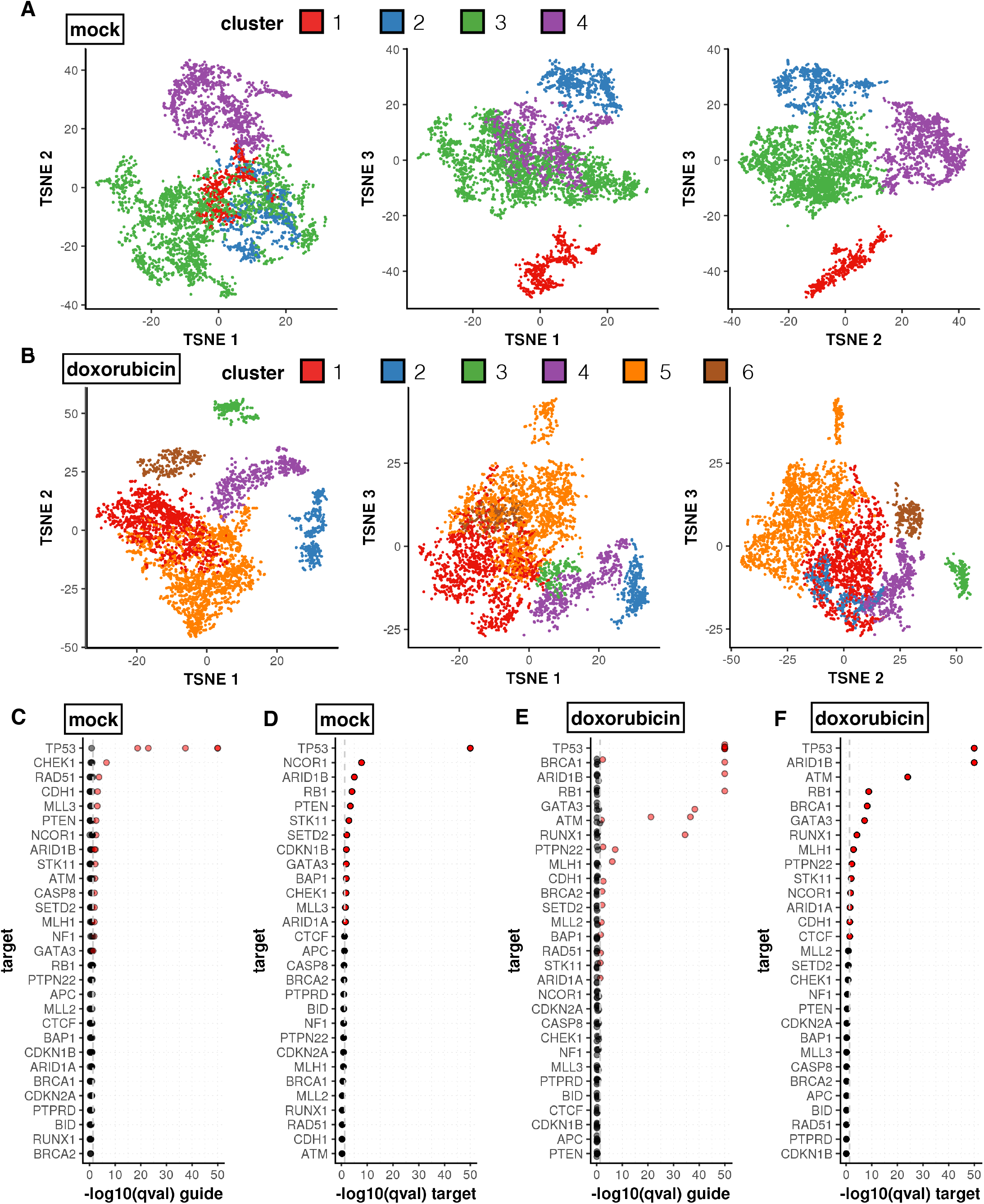
Loss of several targets alter the distribution of mock and doxorubicin exposed cells within tSNE clusters. **A-B)** 3D tSNE embedding and clustering of mock and doxorubicin treated samples, respectively. **C-F)** Chi-squared test qvalues (p values adjusted using the Benjamini-Hochberg method) resulting from testing for differences in the distribution of targets in our screen at both the individual sgRNA **(C and E)** and overall target levels **(D and F).** Comparisons are relative to the distribution of non-targeting controls across tSNE clusters for mock and doxorubicin treated samples, respectively (qvalues were capped to 1e-50 for visualization). Significant differences below a qvalue of 0.05 are colored in red (boundary marked by the grey dashed line).

**Figure S8.**
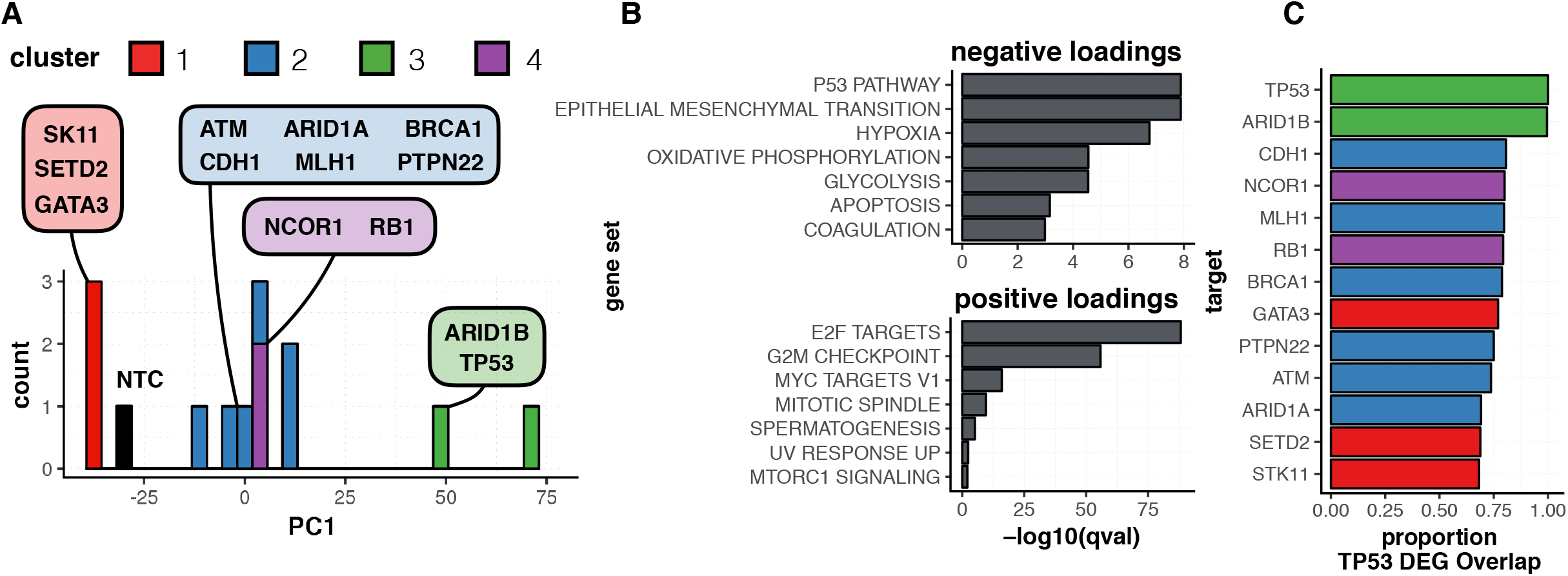
Enriched target-cluster pairs highlight tumor suppressors that share various degrees of a *TP53* deficient signature **A)** Fisher's exact with weights applied to guides according to an expectation maximization procedure were performed for the doxorubicin treated sample to find clusters from Fig. S7 panel B were particular targets were found to be enriched. Cells with target-cluster pairs that showed enrichment were used to generate an aggregate expression profile for every target within genes that are differentially expressed between *TP53* and non targeting controls (NTC). A PCA was performed on these average expression profiles and a distribution of targets across PC1 is shown colored by the cluster in which they were found to be enriched. **B)** Gene set enrichments for top positively and negatively loaded (less than -0.02 or greater than 0.02) genes along PC1 (qval < 0.01). **C)** Differential expression tests were performed for cells within each enriched target-cluster pair, comparing each target to all NTC cells. The proportion of overlap between these differentially expressed genes and the genes differentially expressed between *TP53* and NTC is shown.

**Figure S9.**
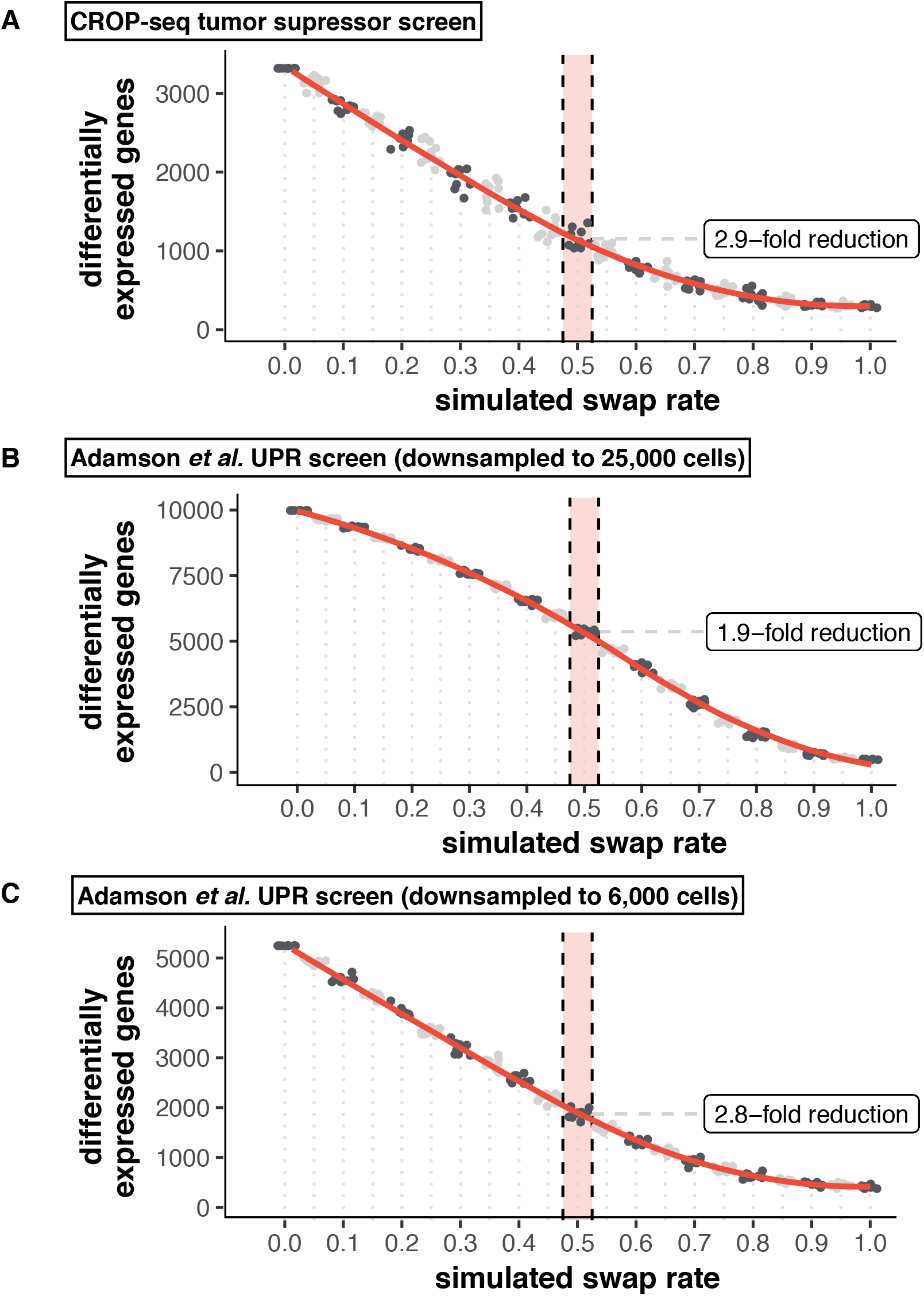
Swap rate simulations for our own CROP-seq tumor suppressor screen and the unfolded protein response screen from Adamson *et al*. Each dataset was subjected to simulation of progressively higher fractions of target assignment swapping to mimic the impact of template switching. Number of differentially expressed genes across the target label at FDR of 5% is plotted at each swap rate. 0.5 corresponds to the 50% swap rate determined via FACS. **A)** CROP-seq tumor suppressor screen from our study. **B)** Unfolded protein response screen from Adamson et al. downsampled from ∼50,000 to 25,000 cells to make simulations computationally feasible. **C)** Unfolded protein response screen from Adamson *et al*. downsampled to 6,000 cells to illustrate how reduced power impacts the observed impact from simulated swapping.

